# Semantic Networks Generated from Early Linguistic Input

**DOI:** 10.1101/157701

**Authors:** Andrei Amatuni, Elika Bergelson

**Author notes:** {, }.

## Abstract

Semantic networks generated from different word corpora show common structural characteristics, including high degrees of clustering, short average path lengths, and scale free degree distributions. Previous research has disagreed about whether these features emerge from internally or externally driven properties (i.e. words already in the lexicon vs. regularities in the external world), mapping onto preferential attachment and preferential acquisition accounts, respectively (Steyvers & Tenenbaum, 2005; Hills, Maouene, Maouene, Sheya, & Smith, 2009). Such accounts suggest that inherent semantic structure shapes new lexical growth. Here we extend previous work by creating semantic networks using the SEEDLingS corpus, a newly collected corpus of linguistic input to infants. Using a recently developed LSA-like approach (GLoVe vectors), we confirm the presence of previously reported structural characteristics, but only in certain ranges of semantic similarity space. Our results confirm the robustness of certain aspects of network organization, and provide novel evidence in support of preferential acquisition accounts.

## Introduction

A word functions as an atomic unit of meaning, in principle carrying independent semantic content. In practice though, it occurs with its fellow words, as humans produce language. From this word-stream, infants begin to understand words by 6-9 months (Bergelson & Swingley, 2012), and to produce them soon thereafter. Here we aim to shed light on how these semantic atoms are organized in the mental lexicon, and the degree to which this representational structure is reflective of the conceptual order found “out there” in the world.

To explore this, we turn to semantic networks, an idea dating back nearly a century (Trier, 1931). Given that words are related along semantic dimensions, characterizing these relations is a first step towards understanding their representational structure. Previous research on semantic networks generated from word corpora have shown small-world connectivity (i.e. any given word node is not very many nodes away from any other), as well as scale free degree distributions (i.e. a few nodes serve as ‘hubs’, and node distribution follows a power law such that *probability(k)* ≈ k^−α^, for a node with degree *k,* and scaling parameter α) (Sigman & Cecchi, 2002; Steyvers & Tenenbaum, 2005; Hills et al., 2009). This suggests that semantic information may be inherently structured in nonrandom, clustered, and highly organized ways, which internal representations may mirror or exploit^1^ (Todd, Hills, & Robbins, 2012). Scale invariance (here equivalent to scale‐free distributions), has been found in many cognitive domains and diverse natural phenomena; it is argued to be a general unifying principle of cognitive organization (Kello et al., 2010).

Barabási and Albert (1999) suggest that graphs with degree distributions that follow power laws imply constraints on the processes which formed them. Their model for generating such networks relies on incremental growth and a process of “preferential **attachment**” (hereafter PAT), whereby existing nodes with many connections are preferentially “chosen” by new nodes. While their resulting graphs display power law degree distributions, they did not find small world connectivity of the kind found in semantic networks, such as those generated from WordNet and Roget’s Thesaurus. Building on this, Steyvers and Tenenbaum (2005) proposed a model for incrementally growing semantic networks similar to Barabási and Albert (1999), which indeed resulted in both small world and scale free structures. Their growth process centered on semantic differentiation, i.e. new words that are more *contrastive* with existing words are preferentially incorporated into the graph; they include a frequency parameter as well. The resulting semantic graphs showed degree distributions which reflected the relative time at which a particular node was added to the network: age-of-acquisition (AoA) norms for words corresponded to the relative number of connections in these graphs. PAT-based graphs inherently bias nodes which are added earlier to have higher degree.

Steyvers and Tenenbaum (2005) suggest that the structure of *internal* representations guides the selection process of new words or concepts. In contrast, Hills et al. (2009) propose that the connectivity of words in the *external* environment plays a guiding role in the acquisition of new words. In this alternative, dubbed preferential **acquisition** (hereafter PAQ), the relative salience between unlearned words directs new node integration into the lexicon. Under PAQ, the structure of the *external semantic ground* is itself scale-free, clustered, and small world, leading internal representations to mirror this structure as lexical items are added. This contrasts with PAT, which suggests that the structuring is a consequence of incremental semantic network growth. Under PAQ, the higher a word’s contextual variety, the more interactions it has with other elements in the external ground. This results in more neighbors in semantic network space, making it more linguistically salient to the learner. Indeed, evidence by Hills, Maouene, Riordan, and Smith (2010) suggests a role for contextual variety and associative density in *noun* lexical development in particular. In the present work, we build on these previous results, combining approaches that suggest network properties arise from *incremental* generative processes with networks that are definitively *non‐generative,* as a window into how external and internal semantic spaces may influence the growing noun lexicon.

One limitation of previous work concerns operationalizing semantic relatedness, generally achieved through hand tagged features or word associations (Steyvers & Tenenbaum, 2005; Hills et al., 2009). Hand-tagged features may not reflect the underlying semantic organization given that they stem from an overt metalinguistic task. Indeed such features do not produce scale free graphs or predict AoA. Word association data lead to directed networks, which may obscure inherent transitivity between word pairs (unless both words have the other as their associative target). It’s not clear whether this directedness is inherent in the lexicon. While asymmetry is conspicuous in human similarity judgments (Tversky, 1977), this may be a function of task rather than underlying semantic representations. In the present work, we rely on neither hand-tagged features nor directed associations in building our semantic graphs.

Given the goal of explaining how semantic structure emerges, a further limitation of previous work lies in the constituent word nodes in the semantic networks, which used e.g. free associations, Roget’s Thesaurus, and WordNet, rather than child-directed corpora. In this paper, we make use of a new corpus of words from infant‐caretaker interactions. This allows us to examine whether scale free distributions, small world connectivity, and links to lexical development trajectories are limited by corpus origin, and thus whether using a full range of concrete nouns children are exposed to in naturalistic settings renders different results.

Here we extend previous work and begin to address these limitations by building ecologically valid semantic graphs of early linguistic input. We use modern vector space methods to calculate undirected semantic relations, resulting in a gradient of networks parameterized by degree of similarity. We limit network nodes to only those which infants’ hear and embed them in a space which approximates a common semantic ground shared by infants and adults alike. We also investigate links between word frequency in the corpus, and connectivity rank in network space.

## Present Study

We generate networks using a new model of semantic relatedness: vectors trained with GloVe (Pennington, Socher, & Manning, 2014). We first determine whether our networks reproduce previously reported small world structure, and scale invariance (i.e. power law distributions). Such structures are consistent with PAT or PAQ. However, only PAT proposes that such structures arise due to incremental growth mechanisms (Barabási & Albert, 1999). PAT suggests that words already in the *internal lexicon* guide new word selection: early words have higher degree than later-added words, i.e. new additions “prefer” to attach to words with higher degree. In contrast, PAQ proposes that external network connectivity drives node addition, suggesting that internal structures mirrors external structure, which may be scale-free, small-world, or not. Because our networks are built using the GloVe vectors, they are, by definition, non-generative and non-incremental: showing scale free and small world behavior in our networks would suggest this structuring might exist without PAT’s assumed incremental generative growth processes.

As a proxy for AoA, we make use of parent-reported vocabulary norms from WordBank (Frank, Braginsky, Yurovsky, & Marchman, 2016), a compilation of the MacArthur-Bates Communicative Developmental Inventory (CDI.) We assume words known by more infants at a given age have been in the lexicon longer. Here we attempt to replicate network structure and AoA correlations originally presented as evidence for PAT, while violating PATs assumption of incremental growth. If successful, it would imply that scale-free structure does not itself depend on PAT.

We also test for evidence of PAQ, by determining whether words that go from being poorly-known to well-known over time have more connections in the externally-based network than those that remain poorly known over time. That is, we test PAQ’s proposal that high degree nodes in networks generated from external linguistic input are acquired *earlier* than lower degree nodes in those same networks. Notably, PAQ models do not depend on power law distributions or scale free behavior, but rather on children selectively integrating *salient* (more densely connected) words from all possible lexical items they’re exposed to. If adult sampling is also inherently biased to those words which have high degree in semantic network space, then we expect too that highly frequent words have higher connectivity relative to all child-directed words. Because our corpus is generated from a large sample of child-directed speech, we can further compare word frequency statistics with degree distributions generated using the same set of words.

## Method

### Data

The SEEDLingS corpus (Bergelson, 2016a, 2016b) comes from home recordings of 44 infants from upstate New York, followed from 6 to 17 months. Each month, a daylong audio recording and hour-long video recording were collected. All videos and 3-10 hours of each audio recording were manually tagged for concrete nouns directed to and/or attended by the child, creating tags of several thousand hours of naturalistic interactions between infants and caregivers. We exclude utterances made by the child, resulting in a final dataset of 4359 unique noun-types (194204 tokens). Plurals and diminutives were consolidated into a “basic level” proxy for word lemmas for each recording. These nouns were used to generate the **SEEDLingS-All** graphs. We also generate graphs for 6 month recordings alone (1855 types, 29289 tokens; **SEEDLingS-6mo**) and a 16/17 month combined set (1708 types, 26969 tokens; **SEEDLingS-16+17mo**), to contrast networks generated from speech to pre-verbal infants and speech to newly verbal toddlers; the SEEDLingS networks are our model of the *external* linguistic environment. We generated an additional network (**WordBank**), using only the 369 nouns on the CDI; this serves as our *internal* semantic network, given that it only includes words that (some) 16-30- month-olds produce.

As our measure of relative AoA, we used by-word summary data from the online WordBank repositories (Frank et al., 2016). This data includes productive vocabularies for children aged 16 to 30 months (reported productive vocabulary is generally more reliable than reported receptive vocabulary.^2^ We make the assumption that words said by more children at a given age entered the lexicon earlier. Indeed, the age at which a word is produced by 50% of children, the AoA metric used by Hills et al. (2010), is significantly inversely correlated with the percentage of production at 16,23, and 30 months (r = −0.78, −0.97, and −0.88 respectively; all *p <* 0.005). Furthermore, AoA measures correlate with children’s elicited naming rates (Morrison, Chappell, & Ellis, 1997). We use WordBank norms rather than the SEEDLingS infants’ own productions, as an independent and extremely large-n (n = 5450, for English) estimate of children’s knowledge for each word in our networks, removing potential dependencies in our analyses.

### GloVe Vectors

Since our dataset is not tagged with semantic features, and since results with hand-engineered features have been mixed, we chose to follow a method described by Steyvers and Tenenbaum (2005) and use semantic vector space models to generate edges between any nodes above a given similarity threshold. We build on their use of Latent Semantic Analysis (LSA) vectors. In that work, LSA vectors (which along with other geometric methods, are non-incremental) did not generate scale free networks; this result was used to suggest that such approaches are incompatible with incremental growth and PAT. To generate our graphs, we use pre-trained word vectors produced by GloVe, a recently developed algorithm for word embedding (Pennington et al., 2014). Using this algorithm, we can investigate whether we find scale free and small world graphs; if so, the original failure to do so might be LSA-specific, and *not* a necessary consequence of PAT, as the authors suggested.

GloVe has been demonstrated to have higher performance on many different word similarity tasks compared to word2vec and matrix factorization methods using SVD. Here, we opted to use vectors trained on the Common Crawl corpus with 42 billion tokens, resulting in 300 dimensional vectors for 1.9 million unique words.^3^ In some sense this ‘full’ dataset provides word similarity proxy based on the *target* (i.e. adult) meanings the child is acquiring. Further analyses using vectors trained on CHILDES (MacWhinney, 2000), displayed analogous and in some cases even stronger patterns than the current results.^4^ This to us suggests consistency in the linguistic manifestation of word meaning (and perhaps their concomitant cognitive processes) at both large and narrow-sampling scales.

Similar to LSA, GloVe learns vector representations of words from co-occurrence matrices built from large text corpora. It instantiates the distributional hypothesis of linguistics, famously articulated by Firth (1957): “you shall know a word by the company it keeps”. Because the GloVe vectors encode co-occurrence statistics derived from natural language, our similarity measures also indicate the degree to which two words share contextual coherence. I.e., the more connections a word has in the semantic network, the more words it shares this coherence with. Given this high dimensional encoding space, we can use a continuous metric of similarity. Iterating through similarity thresholds, we create a gradient of networks to study.

### Generating Semantic Networks

We generate graphs across a range of similarity thresholds (ε). Our similarity measure is the cosine between two GloVe vectors. The cosine function also normalizes for word frequency (to some degree) since dot products are divided by their vector norms. For each corpus, for each word, we calculate cos(θ) between it and every other word in the set.

We give an undirected edge between two words if their cosine is above a threshold ε. Since generating each graph is a quadratic operation we normalize the vectors to unit length before calculating cosines. We iterate ε from 0 to 0.99 (step size=0.01), generating a graph for each similarity threshold. Further methods of edge generation are left for future research. Our code and IPython notebooks are on Github^5,6^.

## Results and Discussion

### Correlations Between Node Degree and Production

We generated 100 graphs for each corpus, one for each value of ε. We calculated Spearman’s rank correlation coefficients between each word’s number of connections and productive vocabulary norm (for the 369 CDI nouns), for each network and similarity threshold, at 16, 23 and 30 months. Under both PAT and PAQ, we would expect to see that words with more connectivity have higher CDI production rates. Indeed, we find robust and significant correlations between the degree of a word in the network, and the percent of toddlers who produced it, for a range of ε, across corpora and ages; Fig.1.

**Figure 1:**
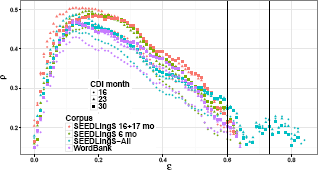
Correlation coefficients (p) between number of edges and CDI production rate across words, as a function of similarity threshold ε (all p are significant at *p <* 0.005). Color indicates which corpus created the network, shape indicates which month of CDI norms was used to calculate p. Vertical lines indicate the range in which we find scale free degree distributions (0.6-0.73). (For thresholds ε > 0.75 there were very few (or no) edges being created, explaining the discontinuity and lack of points towards the end of the scale.)

More specifically, we find similar behavior across all networks, with a global peak in correlation for ε = 0.12-0.19. All peak correlation values had Spearman’s ρ = 0.43-0.52, with *p <* 10^−5^, showing consistent behavior across networks and ages. This suggests that both the parent’s word choice given a child’s age, and the child’s responsiveness to external semantic density across time are roughly constant. This range of ε where the correlation is at a maximum is relatively low, allowing very loose semantic associations to result in edges. In Figure 2 we show a subgraph from the SEEDLingS-All network, centered around the node “baby.”

**Figure 2:**
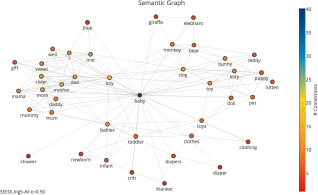
“Baby” subgraph from the SEEDLingS-All network at the similarity midpoint (ε = .5), where “baby” has 40 neighbors.

### Power Law Degree Distributions

Surprisingly, at the values of ε where the correlation is maximal, we did not find power law distributions. We did however find them at higher thresholds: at ε = 0.68 we can fit a power law function with α = 3.2 *±* 0.1, with a log likelihood ratio in favor of power law over exponential fit (*R* = 115.73, *p* = 2.337 × 10^—21^). Indeed, at ε = 0.63 — 0.75 we find power law distributions (α =2.39-3.73) characteristic of scale free networks. At these higher thresholds a word’s neighbors are semantically very close, similar to other semantic graphs which have shown scale free distributions (e.g. Roget’s Thesaurus), suggesting this property might depend on connections’ high semantic proximity. See Figure 3.

**Figure 3:**
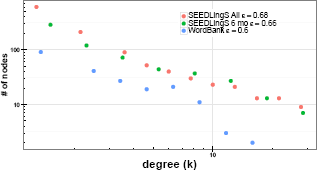
Sample networks showing degree distributions with power law behavior (α=3.1-3.2; SE=.1, *p <* 0.001; similar behavior found across networks for 0.60 ≤ ε ≤ 0.73). Distributions are plotted on a log-log scale with logarithmic spacing between points, which represent the edges of bins. Power law distributions appear linear on this scale.

PAT models presuppose power law distributions (indeed, PAT was initially proposed after observing scale free distributions in semantic networks, and arguing that this limits the kinds of mechanism which could have created them). We thus further analyze the range of ε where our networks display power law behavior. Again, networks showing this distribution are critical for PAT (and thus our CDI-based Wordbank networks’ proxie of an *internal* network), but incidental for PAQ, which makes no claims about power-law distributions.

Limiting our focus to these ranges (between the vertical lines in Figure 1,) we see that the degree of a given word in the SEEDLingS-All network has uniformly higher correlation with productive vocabulary norms compared to that same node in the internal (i.e. WordBank) networks. (To be clear, we can only calculate ρ for words we have CDI norms for, but the SEEDLingS networks contain all the nouns infants heard, while the WordBank networks contain *only* CDI word nodes). This pattern is consistent with PAQ, where more densely connected words in the environment are preferentially incorporated into the learner’s lexicon. The correlation between AoA and node degree for the WordBank networks, along with their scale-free organization, suggest that PAT is *not* a necessary condition for this behavior, since these graphs were generated using GloVe. The presence of these same correlations for our other networks (which serve as a proxy of an externally-based network) in this same range of ε, are also scale-free, and provide new support for PAQ.

This pattern validates our method of generating graphs using GloVe vectors: both the WordBank and our SEEDLingS networks display behavior consistent with previous accounts (i.e. scale free distributions in internal lexical networks and node degrees correlated with AoA). If anything, the SEEDLingS networks show the predicted structure more strongly, suggesting the nouns infants actually hear may form a better representation than limiting the space to lexically simple, early-learned nouns alone. Our current analysis suggests that scale-free and small world structure can be produced *without* an incremental growth process, since our graphs were generated using a vector space model (i.e. GloVe). If words are to be incorporated using PAQ, this external structure would necessarily be mirrored in the internal lexical network. However, it remains possible that PAT and PAQ could both be at play during infant lexical development, perhaps with PAQ supplementing PAT by providing a structured sampling space for new word selection.

### Clustering Coefficients and Path Lengths

We next examined whether our semantic networks, based on natural language to children, exhibited two key small-world properties found in previous research: low average path length (L), and high clustering coefficients (C). We found that the SEEDLingS networks generally had lower C and higher L than the WordBank networks. In Table 1 we list C and L of each network at their respective peak from Figure 1. We also generated Erdős-Rényi and Watts-Strogratz graphs for comparison (Watts & Strogatz, 1998; Erdős & Rényi, 1960); Erdős-Rényi gives us a baseline measure of a comparably sized graph built using a *random* process, while Watts Strogatz provides a prototypical example of a small world graph, with low L and high C. The SEEDLingS networks clearly showed higher clustering coefficients and smaller average path lengths compared to the Erdős-Rényi graph, and comparable behavior to the Watts-Strogatz graph. This small world organization is indicative of hub structures in the network, where a few very densely connected nodes establish routes between a large proportion of the graph, keeping the average shortest path length low. This is also a defining feature of networks with power law distributions, even though the networks we’ve listed in Table 1 do not fit that criteria. This small world organization, even in the absence of power law distributions,^7^ supports previous findings in other semantic networks and suggest that even in child-directed natural language input we see these structures.

**Table 1:** Clustering coefficients (C) and average shortest path lengths (L) of the largest connected subgraph at peak values from Figure 1. Generated Erdős-Rényi (n=6404, p=0.05) and Watts-Strogratz graphs (n=6404, k=64, p=0.05) are listed for comparison.

| Corpus | $\epsilon$ | C | L |
| --- | --- | --- | --- |
| SEEDLingS All | 0.13 | 0.594 | 1.749 |
| SEEDLingS 6 mo. | 0.16 | 0.669 | 1.739 |
| SEEDLingS 16+17 mo. | 0.12 | 0.726 | 1.534 |
| WordBank | 0.13 | 0.895 | 1.202 |
| Erdős-Renyi | - | 0.049 | 1.950 |
| Watts-Strogatz | - | 0.634 | 3.013 |

### AoA as a Function of Frequency and Connectivity

The SEEDLingS corpus contains word frequency counts, a particularly powerful predictor of word acquisition (Goodman, Dale, & Li, 2008), allowing us to examine the relationship between word frequency and network connectivity.^8^A positive relationship would suggest that densely connected words are preferentially sampled in adult speech directed towards children. As shown above, more highly connected words were said by more toddlers, across our networks (at peak e, all p > .51, all *p <* .0001; see Fig. 1). Using model comparison of simple linear models, we find that including both word frequency and node degree as predictors of word production (at 16, 23, and 30 months) accounts for significantly more variance than either alone (all *p <* .01; interaction term significantly improved model fit for months 23 and 30 only, both *p <* .01).^9^

Finally, to better understand whether our semantic networks find support for PAT and PAQ models, we tested one specific prediction of each. For PAT, we tested whether words that had been known longer (i.e. by proxy, were said by more children) had more connections than those that had not. Indeed, conducting a median split on words’ production rates at each age (16, 23, and 30 months), we find that better known words have higher degree than less well known words (p < 0.005 by Wilcoxon Test). For PAQ, we tested whether the words that went from less-well-known to better-known over 16-30 months had higher degree than those that remained poorly known. Indeed, for words produced below the median rate at 16 months, those *below* the median at 30 months had significantly lower degree than those above it (p < .005 by Wilcoxon Test.) This supports PAQ’s proposal that high degree nodes in networks generated from *external* linguistic input are acquired earlier than lower degree nodes in those same networks.

## Conclusions

Our results suggest there is inherent semantic structure present in the early linguistic environment, and that both the caregivers and their children are likely sensitive to this nonuniform distribution of semantic information. Because the SEEDLingS corpus provides a uniquely rich dataset of early linguistic input, we were able to construct ecologically valid networks and study differences in their structure across time for a constant set of infants. Our present findings support previous work addressing semantic network structure. Using a modern semantic vector space model to generate our graphs, we were able to confirm the presence of scale free degree distributions in our networks, as well as high clustering coefficients and low average path lengths. This method for generating semantic networks avoids the need for hand engineered features and sidesteps the limits of free-association data, providing a potentially more advanced measure of semantic relatedness compared to the original work on PAQ.

That said, the process that generated the GloVe vectors here is not the same as that generating any human’s lexicon; further work is needed to strengthen and test links between these representations. Moreover, the GloVe model does not speak to the *origin* of token distributions in natural language. It does, however, encode a geometric projection of a meshwork of causal substructures present in the external world. Future research will explore the link between these structures and their grounding in cognitive processes. While we have taken a few steps towards examining network growth over time (finding little difference in our 6mo. and 16+17mo. SEEDLingS networks, or over 16, 23, and 30mo. CDI norms), more work is needed to better understand not only *whether* PAT and/or PAQ-compatible processes are at play, but *how* the interplay between input and uptake changes as the learner grows.

In their original work, Hills et al. (2009) were not able to produce scale free graphs using their hand made features, but were able to do so using adult free association data. In our own graphs we saw that the scale free property only manifested at relatively high values of ε, where only very closely related words (often synonyms) were connecting to each other. Because our measure of similarity was parameterized, we were able to produce a gradient of networks and study their behavior across a range of thresholds, focusing at different ranges of the scale as needed. By generating scale free networks using a non-incremental procedure, we lend support to the hypothesis that this structuring may be an inherent feature in the external environment, rather than a consequence of how it’s integrated into internal representations. Building on Firth: our results suggest that words may indeed *become* known by the company they keep, and that the relevant neighbors may be both those inside the lexicon, and in the as-yet unknown external world of words.

## Acknowledgments

We thank the SEEDLingS team, and NIH DP5-0D019812.

## Footnotes

1 Graphs with high clustering coefficients and low average path lengths, as in small-world networks, are efficient to search and relay information through, while scale invariance allows a single algorithm to operate across seemingly disparate representational frames.

2 SEEDLingS networks contain many more nouns than the WordBank network (resulting in different connectivity patterns), but AoA data is only available for the 369 CDI nouns for all networks

3 http://nlp.stanford.edu/projects/glove/

4 We omit these due to space, but thank an anonymous reviewer for this suggestion; they will be presented at CogSci.

5 https://github.com/andreiamatuni/wordgraph

6 https://github.com/BergelsonLab/semspace

7 Scale free networks are inherently ultrasmall (Cohen & Havlin,2003)

8 Phonological neighborhood effects are saved for future work

9 Node degrees are from the SEEDLings-All network at peak similarity threshold of e = 0:13

